# Proteomic fingerprint identification of Neotropical hard tick species (*Acari: Ixodidae*) using a self-curated mass spectra reference library

**DOI:** 10.1101/2020.04.13.040089

**Authors:** Rolando A. Gittens, Alejandro Almanza, Eric Álvarez, Kelly L. Bennett, Luis C. Mejía, Javier E. Sanchez-Galan, Fernando Merchan, Jonathan Kern, Matthew J. Miller, Helen J. Esser, Robert Hwang, May Dong, Luis F. De León, Jose R. Loaiza

**Affiliations:** Centro de Biodiversidad y Descubrimiento de Drogas, Instituto de Investigaciones Científicas y Servicios de Alta Tecnología (INDICASAT AIP), Panama, Republic of Panama; Centro de Neurociencias, INDICASAT AIP, Panana, Republic of Panama; Smithsonian Tropical Research Institute, Panama, Republic of Panama; Grupo de Investigación en Biotecnología, Bioinformática y Biología de Sistemas, Facultad de Ingeniería de Sistemas Computacionales, Universidad Tecnológica de Panamá, Panama, Republic of Panama; Grupo de Investigación en Sistemas de Comunicaciones Digitales Avanzados, Facultad de Ingeniería Eléctrica, Universidad Tecnológica de Panamá, Panama, Republic of Panama; ENSEIRB-MATMECA – Bordeaux INP, France; Sam Noble Oklahoma Museum of Natural History and Department of Biology, University of Oklahoma, Norman, OK, USA; Department of Environmental Sciences, Wageningen University, Wageningen, the Netherlands; Department of Biology, Swarthmore College, Swarthmore, PA, USA; Department of Biology, University of Massachusetts Boston. Boston, MA, USA; Programa Centroamericano de Maestría en Entomología, Universidad de Panamá, Panama, Republic of Panama

## Abstract

Matrix-assisted laser desorption/ionization (MALDI) time-of-flight mass spectrometry is an analytical method that detects macromolecules that can be used as biomarkers for taxonomic identification in arthropods. The conventional MALDI approach uses fresh laboratory-reared arthropod specimens to build a reference mass spectra library with high-quality standards required to achieve reliable identification. However, this may not be possible to accomplish in some arthropod groups that are difficult to rear under laboratory conditions, or for which only alcohol preserved samples are available. Here, we generated MALDI mass spectra of highly abundant proteins from the legs of 18 Neotropical species of adult field-collected hard ticks, several of which had not been analyzed by mass spectrometry before. We then used their mass spectra as fingerprints to identify each tick species by applying machine learning and pattern recognition algorithms that combined unsupervised and supervised clustering approaches. Both principal component analysis (PCA) and linear discriminant analysis (LDA) classification algorithms were able to identify spectra from different tick species, with LDA achieving the best performance when applied to field-collected specimens that did have an existing entry in a reference library of arthropod protein spectra. These findings contribute to the growing literature that ascertains mass spectrometry as a rapid and effective method for taxonomic identification of disease vectors, which is the first step to predict and manage arthropod-borne pathogens.

**Author Summary:** Hard ticks (Ixodidae) are external parasites that feed on the blood of almost every species of terrestrial vertebrate on earth, including humans. Due to a complete dependency on blood, both sexes and even immature stages, are capable of transmitting disease agents to their hosts, causing distress and sometimes death. Despite the public health significance of ixodid ticks, accurate species identification remains problematic. Vector species identification is core to developing effective vector control schemes. Herein, we provide the first report of MALDI identification of several species of field-collected Neotropical tick specimens preserved in ethanol for up to four years. Our methodology shows that identification does not depend on a commercial reference library of lab-reared samples, but with the help of machine learning it can rely on a self-curated reference library. In addition, our approach offers greater accuracy and lower cost per sample than conventional and modern identification approaches such as morphology and molecular barcoding.

## Introduction

Hard ticks (Ixodidae) are hematophagous ectoparasites that feed on almost every species of terrestrial vertebrate on earth, including *Homo sapiens sapiens* [1, 2]. Due to a complete dependency on blood as a food source, both sexes of adults and immature ticks are capable of transmitting disease pathogens to their hosts, causing significant morbidity and sometimes even death [3, 4]. Research on hard ticks has increased recently in the Neotropics, where a growing number of outbreaks of tick-borne related illnesses have been documented [5–8]. Despite these efforts, comprehensive studies about the ecology, behavior and control of hard ticks relevant to public health remain elusive in Central America due to the shortcomings of traditional taxonomic methods to species identification. Taxonomic identification of Neotropical Ixodidae has traditionally relied on adult morphological characters [9]; however, morphological keys for immature stages (i.e., larvae and nymphs) are lacking and experts are often unable to reliably identify immature ticks to species [9, 10]. Moreover, morphological identification of ticks is unrealistic in epidemiological settings because assessing the role of ticks as disease vectors usually involves identifying hundreds of individuals for pathogen screening, an extremely time-consuming effort, which may be further impeded by the lack of qualified taxonomic specialists [11].

Matrix-assisted laser desorption/ionization (MALDI) time-of-flight mass spectrometry is an analytical technique that allows for sensitive and accurate detection of complex molecules such as proteins, peptides, lipids and nucleic acids [12–14]. The conventional MALDI approach has been used successfully to generate markers for proteomic identification of microorganisms such as pathogenic bacteria and fungi, which can be cultured in the laboratory and form discrete colonies with very consistent mass spectra that facilitates the development of reference libraries for identification of unknown samples [15, 16]. In fact, a commercial program offered by the manufacturers of the MALDI technology is capable of determining statistical similarities between the spectra of unknown samples and a well-curated, proprietary reference library of bacteria and fungi to identify the species of the unknown specimen. This is analogous to the process of matching fingerprints, and offers a simplified comparison score that ranges from 0.0 to 3.0. Scores above or equal to 2.3 represent a confident match at the genus rank, and high probability at the species level, while values below 1.7 are considered as non-reliable identifications [15–17].

Although more challenging than identifying bacteria and fungi due to the size and heterogenicity of the specimen, MALDI has also been used to discriminate among species of invertebrates, including mosquitoes (Culicidae - *Anopheles*), fleas (Pulicidae - Ctenocephalide), biting midges (Ceratopogonidae – *Culicoides*), sandflies (Psychodidae – *Phlebotomus, Lutzomyia*) and ticks (Ixodidae – *Rhipicephalus*) [18–26]. A key finding from these studies is that protein spectra obtained from body sections or whole specimens were similar among individuals of the same morphological species but differed noticeably across different species. Therefore, MALDI protein spectra can be used as a tool to delimit species boundaries in arthropods that are vectors of pathogens. Nevertheless, fresh laboratory-reared specimens are routinely needed to build a reference library that meets the high-quality standards required for classification. This represents an important limitation for some arthropod groups, or assemblages, that are difficult to rear under laboratory conditions. In addition, epidemiological studies often rely on field-collected specimens preserved in ethanol for long-term storage in reference collections. To overcome these limitations, previous studies have opted for adjusting the comparison scores minimum-threshold limit for identification, lowering the manufacturer’s recommended scores from 2.3 to 1.8 [21, 27] or even 1.3 [22, 28]. Hence, mass fingerprinting for the identification of field-collected specimens that do not exist in a reference spectra library (or for those from which reference spectra cannot be generated under ideal conditions) requires an alternative, objective approach [11]. Moreover, most existing applications of MALDI to identify arthropod disease vectors have focused on relatively species-poor vector assemblages from Europe. This technique has been tested less-frequently in the new world tropics [19, 20, 22, 24, 27–36], where vector species richness is the greatest on Earth.

Here, we used MALDI as a scheme to identify Neotropical specimens of adult hard ticks derived from ethanol-preserved field collections. Specifically, we used machine learning and pattern recognition algorithms to classify protein spectra from the legs of field-collected specimens in order to identify a group of unknown samples with a self-curated reference library. MALDI is a promising tool for cataloging and quickly identifying large arthropod groups such as ticks [11]. Our results should contribute to the growing body of literature trying to address questions about feasibility, reliability and universality of the methodology for different environments and species that have not been evaluated before. Properly identifying disease vectors such as Ixodidae in highly diverse Neotropical countries, such as Panama, is a critical first step to predict and manage tick-borne zoonotic pathogens such as *Rickettsia* and arboviruses (e.g., arthropod-borne viruses).

## Methods

### Sample preparation

Ticks stored in ethanol for up to 5 years, and previously identified based on morphological characters, were taken from long-term storage in a −20 °C freezer (S1 Table). A total of 103 specimens from the following species were included in this study: *Amblyomma mixtum (cajennense), Amblyomma calcaratum, Amblyomma dissimile, Amblyomma geayi, Amblyomma nodosum, Amblyomma oblongoguttatum, Amblyomma ovale, Amblyomma pecarium, Amblyomma sabanerae, Amblyomma varium, Amblyomma naponense, Amblyomma tapirellum, Ixodes affinis, Ixodes boliviensis, Dermacentor nitens, Haemaphysalis juxtackochi, Rhipicephalus microplus* and *Rhipicephalus sanguineus*. Samples were prepared following previously published protocols with minor modifications [21, 22]. Briefly, we removed either the left or the right anterior leg from each tick using a scalpel. The leg was then put in tube with 300 μL ultrapure water followed by the addition of 900 μL of 100% ethanol. These tubes were vortexed for 15 s and centrifuged at 13,000 RPM for 2 min. After centrifugation, the supernatant was poured off from the sample tube, which was left to dry for 15 min. Subsequently, the legs were resuspended in 60 μL 70% formic acid and 60 μL 100% acetonitrile and homogenized in the microtube using a manual pestle. The samples were placed in a Branson 1510 ultra-sonicator (Bransonic, Danbury, CT, USA) for 60 minutes in ice water, and then vortexed for 15 s and centrifuged again at 13,000 RPM for 2 min. 1 μL of supernatant was pipetted onto a polished steel MALDI plate and covered with 1 μL of HCCA matrix. After letting the plate dry, it was inserted into the MALDI mass spectrometer to record the protein spectra from tick legs.

### MALDI mass spectrometry parameters

We used an UltrafleXtreme III spectrometer (Bruker Daltonics, Bremen, Germany) to generate the protein mass spectra of each specimen. The equipment has a MALDI source, a time-of-flight (TOF) mass analyzer, and a 2KHhz Smartbeam™-II neodymium-doped yttrium aluminum garnet (Nd:YAG) solid-state laser (λ=355 nm) that we used in positive polarization mode. All spectra were automatically acquired in the range of 2,000 to 20,000 m/z in linear mode for the detection of the most abundant protein ions. Each spectrum represented the accumulation of 5,100 shots with 300 shots taken at a time, and the acquisition was done in random-walk mode with a laser power in the range of 50% to 100% (global laser attenuation at 30%). The software FlexAnalysisTM (Bruker) was used to analyze the mass spectra initially and to evaluate number of ion peaks and their intensity. Visual comparisons of the mass spectra from different tick species gave initial indications of dominant ion peaks that would suggest possible classification into discrete groups. Mass spectra that did not include at least one ion peak with an intensity of 1000 a.u. or more, were considered low quality and filtered out. All samples were placed and measured on three individual target wells with three technical replicates of the mass spectra collected per well.

### Data analysis, clustering algorithms and statistics

The methodology has been described in detail previously by our group for the identification of adult mosquito legs [26], based on similar data analysis for face recognition [37, 38] and spectral classification using mass spectrometry [39, 40]. In brief, 239 mass spectra generated across 103 samples for all 18 species of morphologically-identified Neotropical hard ticks were classified using Principal Component Analysis (PCA) and Linear Discriminant Analysis (LDA), which are linear transformation techniques from the field of

Machine Learning that are commonly used for dimensionality reduction and classification. Dimensionality reduction can help decrease computational costs for classification, as well as avoid overfitting by minimizing the error in parameter estimation.

PCA is an “unsupervised” algorithm that generates vectors that correspond to the direction of maximal variance in the sample space. On the other hand, LDA is a “supervised” algorithm that considers class information to provide a basis that best discriminates the classes (*i.e*., tick species) [37]. For both PCA and LDA analyses, we calculated the Euclidean distance between the vector describing the test sample and the average vector describing each class to identify a test sample. The class with the minimum distance with respect to the test sample was assigned as the identified species for that test sample. The LDA was applied over the data set expressed in terms of the coefficients (*i.e*., principal components) obtained by the PCA. Thus, PCA reduced the dimensionality of the data, and the LDA provided the supervised classification.

The performance of the clustering algorithms was tested using Monte Carlo simulations over 1000 iterations per species to optimize training and cross-validation prediction success rates (Table 2). For each iteration, the data elements in each class were split randomly in approximately, but not less than, 20% of the elements for testing and the rest of the elements for training, for each species. For this analysis, we used the first 150 principal components from the PCA stage that explained 99.9% of the total variance, which after being projected for the LDA algorithm, also generated a 150-components data set. The number of components was chosen after a performance analysis, again using a Monte Carlo approach, that provided the best identification rates. Global and class positive identification rates were calculated to establish the classification capacity of the algorithm (Table 2). The positive identification rate corresponds to the percent ratio between positive identifications performed by the algorithm and the real positive cases in the data.

For visualization purposes in the plots, species that were morphologically identified within the *Rhipicephalus* and *Ixodes* genera were separately compared against *Dermacentor* and *Haemaphysallis* for which there was only one species in each. All species that were morphologically identified within the *Amblyomma* genus were separately compared between themselves or against the *Ixodes* genera.

## Results

Optical micrographs from 18 species of Neotropical hard ticks showed very clear differences among species in terms of adult morphological features (Fig 1, S1 Fig), which was well aligned with the expected unique mass spectra generated from each sample and taxon (Fig 2). The global automatic acquisition rate was 77% for all species (Table 1), confirming that, overall, the mass spectra of field-collected and ethanol preserved specimens allowed automatic acquisition of spectra. In fact, automatic acquisition of spectra results in faster and more objective data acquisition than performing spectra collection manually. The percentage of automatic spectra acquisition with the MALDI ranged from 50 % for *A. mixtum* (*cajennense*), *I. boliviensis* and *R. sanguineus* to 100% for several of the species, including *A. calcaratum, A. geayi, A. sabanerae, I. affinis*, and *R. microplus* (Table 1). The time stored in ethanol or the location of sample origin did not seem to explain the variable percentages of automatic spectra collection (S1 Table). Spectra from freshly collected specimens stored dry at −20 °C, used to establish the methodology, exhibited the best signals, with better-defined spectral peaks and higher signal-to-noise ratio.

In addition, the specimens within each species showed consistently similar protein profiles, regardless of their taxonomic genera, sex, collection date and/or sampling location (Fig 2, Table S1). Mean protein spectra for tick species differed visually among taxa and the differences appeared to be related to their degree of phylogenetic relatedness (Fig 2). For example, species within the genera *Ixodes, Rhipicephalus*, and *Amblyomma* were more similar among themselves in terms of the ions peak number and mass over charge (*m/z*) position in their mass spectra than species from different genera. Nonetheless, some closely related species within the *Amblyomma* genus such as *A. mixtum (cajennense), A. varium*, and *A. tapirellum* also showed fairly distinct protein spectra (Fig 2), which motivated the application of clustering algorithms for their classification.

**Figure 1.**
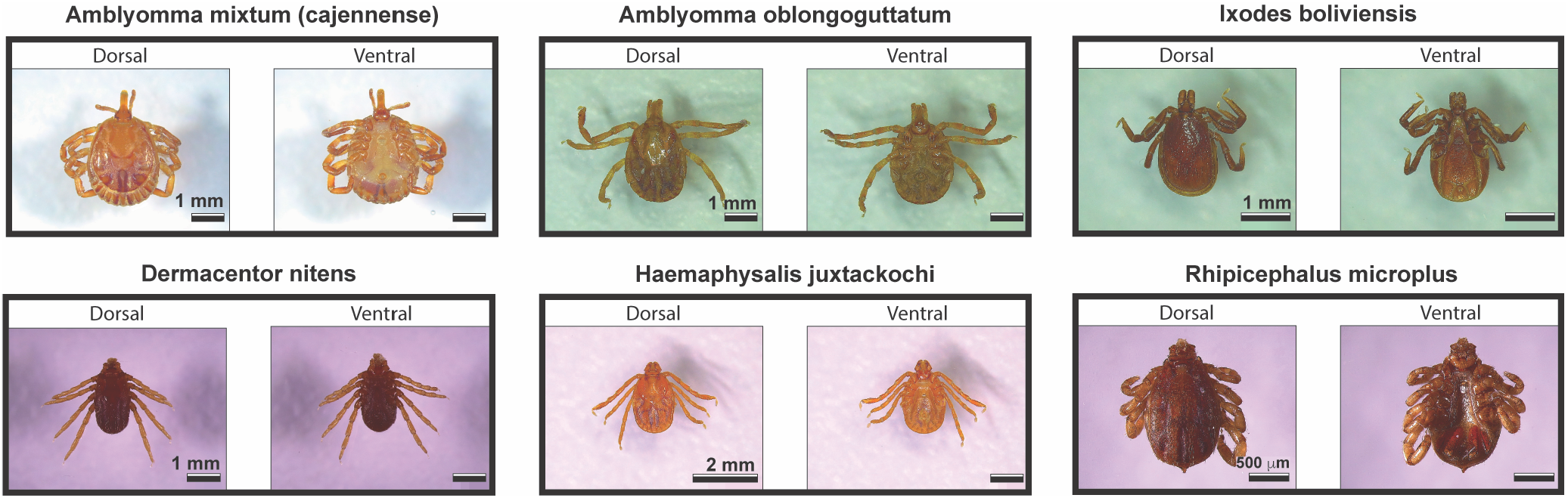
Optical micrographs of Neotropical hard ticks. The image shows the dorsal and ventral sides for 6 of the 18 species of hard ticks in the genus *Amblyomma, Dermacentor, Haemaphysalis, Ixodes* and *Rhipicephalus*. The images for the full assemblage of 18 species can be found in S1 Fig.

**Table 1.** Description of samples subjected to analysis with the MALDI mass spectrometry procedure.

| Species Name | # of samples | Locality code | # of expected spectra | # of obtained spectra | MALDI automatic spectra acquisition rate (%) |
| --- | --- | --- | --- | --- | --- |
| *Amblyomma mixtum (cajennense)* | 4 | a | 12 | 6 | 50% |
| *Amblyomma calcaratum* | 5 | a, b | 15 | 15 | 100% |
| *Amblyomma dissimile* | 4 | c | 12 | 9 | 75% |
| *Amblyomma geayi* | 4 | d | 12 | 12 | 100% |
| *Amblyomma nodosum* | 4 | a | 12 | 10 | 83% |
| *Amblyomma oblongoguttatum* | 4 | a, e | 12 | 8 | 67% |
| *Amblyomma ovale* | 4 | e | 12 | 11 | 92% |
| *Amblyomma pecarium* | 4 | e | 12 | 11 | 92% |
| *Amblyomma sabanerae* | 3 | f | 9 | 9 | 100% |
| *Amblyomma varium* | 4 | g | 12 | 9 | 75% |
| *Amblyomma naponense* | 5 | f | 15 | 9 | 60% |
| *Amblyomma tapirellum* * | 26 | e, g | 78 | 56 | 72% |
| *Ixodes affinis* | 4 | e | 12 | 12 | 100% |
| *Ixodes boliviensis* | 4 | e | 12 | 6 | 50% |
| *Dermacentor nitens* | 4 | c | 12 | 9 | 75% |
| *Haemaphysalis juxtackochi* | 6 | a, e | 18 | 11 | 61% |
| *Rhipicephalus microplus* | 10 | c, d | 30 | 30 | 100% |
| *Rhipicephalus sanguineus* | 4 | a | 12 | 6 | 50% |
| **Total** | **103** | **a-g** | **309** | **239** | **77%** |

(a) = Panama: Chorrera, Las Pavas; (b) = Panama: Colon, Madden Road; (c) = Panama: Colon, Achiote; (d) = Panama: Panama, Capira; (e) Panama: Colon, Barro Colorado Island; (f) Panama: Colon, Sierra Llorona Lodge; (g) Panama: Colon, Gamboa. (\*) Indicates some specific samples that upon collection were stored fresh in Silica Gel (For more metadata information about these samples see also S1 Table).

**Figure 2.**
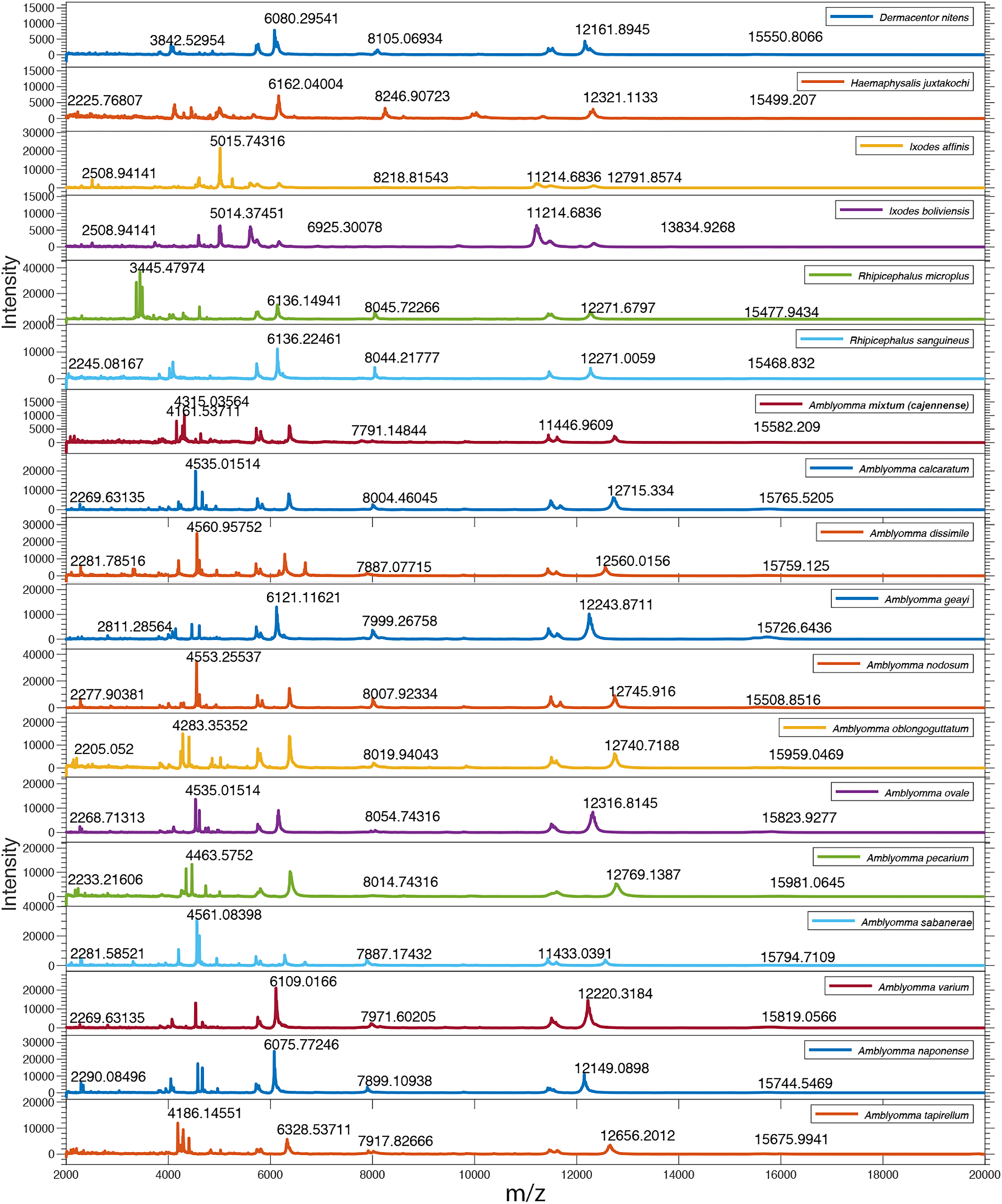
Baseline-corrected and smoothed spectra for 18 species of ticks in the genus *Amblyomma, Dermacentor, Haemaphysalis, Ixodes* and *Rhipicephalus*. Major ion peaks and their molecular weights are annotated in the range of 2,000 to 20,000 m/z for all species.

Distinct mass spectra profiles between morphologically identified ixodid species could be classified by an unsupervised PCA algorithm to identify specimens. The quantitative performance of the PCA algorithm was assessed per species (Table 2), and visually confirmed with the graphic clustering presented in 3D plots (Fig 3). The PCA global positive identification rate was 91.2%, with 14 out of 18 species having higher than 90 % positive identification rate. The PCA graphs showed that most species separated in well-defined clusters, and the distance among clusters seemed to be related to the degree of phylogenetic relatedness as evidenced by the clear separation from the specimens of *Dermacentor* and *Rhipicephalus* with those from *Haemaphysallis* and *Ixodes* (Fig 3A, B), or just between the specimens of *Amblyomma* (Fig 3C). When comparing species within the genus *Amblyomma* against those from *Ixodes*, again the spectra from specimens of each species clustered together with limited overlap between groups and those from different genera were clearly separated (Fig 3D).

**Table 2.** Performance of PCA and LDA clustering algorithms.

| Species Name | PCA Positive<br>Identification<br>Rate (%) | LDA Positive<br>Identification<br>Rate (%) | Spectra<br>per Class | # Training<br>Elements | # Test<br>Elements |
| --- | --- | --- | --- | --- | --- |
| *Amblyomma mixtum*<br>(*cajennense*) | 100.0% | 100.0% | 6 | 4000 | 2000 |
| *Amblyomma calcaratum* | 100.0% | 99.6% | 15 | 12000 | 3000 |
| *Amblyomma dissimile* | 67.6% | 67.6% | 9 | 7000 | 2000 |
| *Amblyomma geayi* | 99.1% | 99.6% | 12 | 9000 | 3000 |
| *Amblyomma nodosum* | 100.0% | 100.0% | 10 | 8000 | 2000 |
| *Amblyomma oblongoguttatum* | 100.0% | 100.0% | 8 | 6000 | 2000 |
| *Amblyomma ovale* | 100.0% | 100.0% | 11 | 8000 | 3000 |
| *Amblyomma pecarium* | 99.8% | 99.0% | 11 | 8000 | 3000 |
| *Amblyomma sabanerae* | 69.3% | 85.9% | 9 | 7000 | 2000 |
| *Amblyomma varium* | 99.8% | 100.0% | 9 | 7000 | 2000 |
| *Amblyomma naponense* | 100.0% | 100.0% | 9 | 7000 | 2000 |
| *Amblyomma tapirellum* | 97.8% | 97.8% | 56 | 44000 | 12000 |
| *Dermacentor nitens* | 21.7% | 45.6% | 12 | 9000 | 3000 |
| *Haemaphysalis juxtackochi* | 90.9% | 97.8% | 6 | 4000 | 2000 |
| *Ixodes affinis* | 84.0% | 89.5% | 9 | 7000 | 2000 |
| *Ixodes boliviensis* | 96.8% | 98.8% | 11 | 8000 | 3000 |
| *Rhipicephalus microplus* | 93.1% | 98.7% | 30 | 24000 | 6000 |
| *Rhipicephalus sanguineus* | 100.0% | 100.0% | 6 | 4000 | 2000 |
| **Global** | **91.2%** | **94.2%** | **239** | **183000** | **56000** |

**Figure 3.**
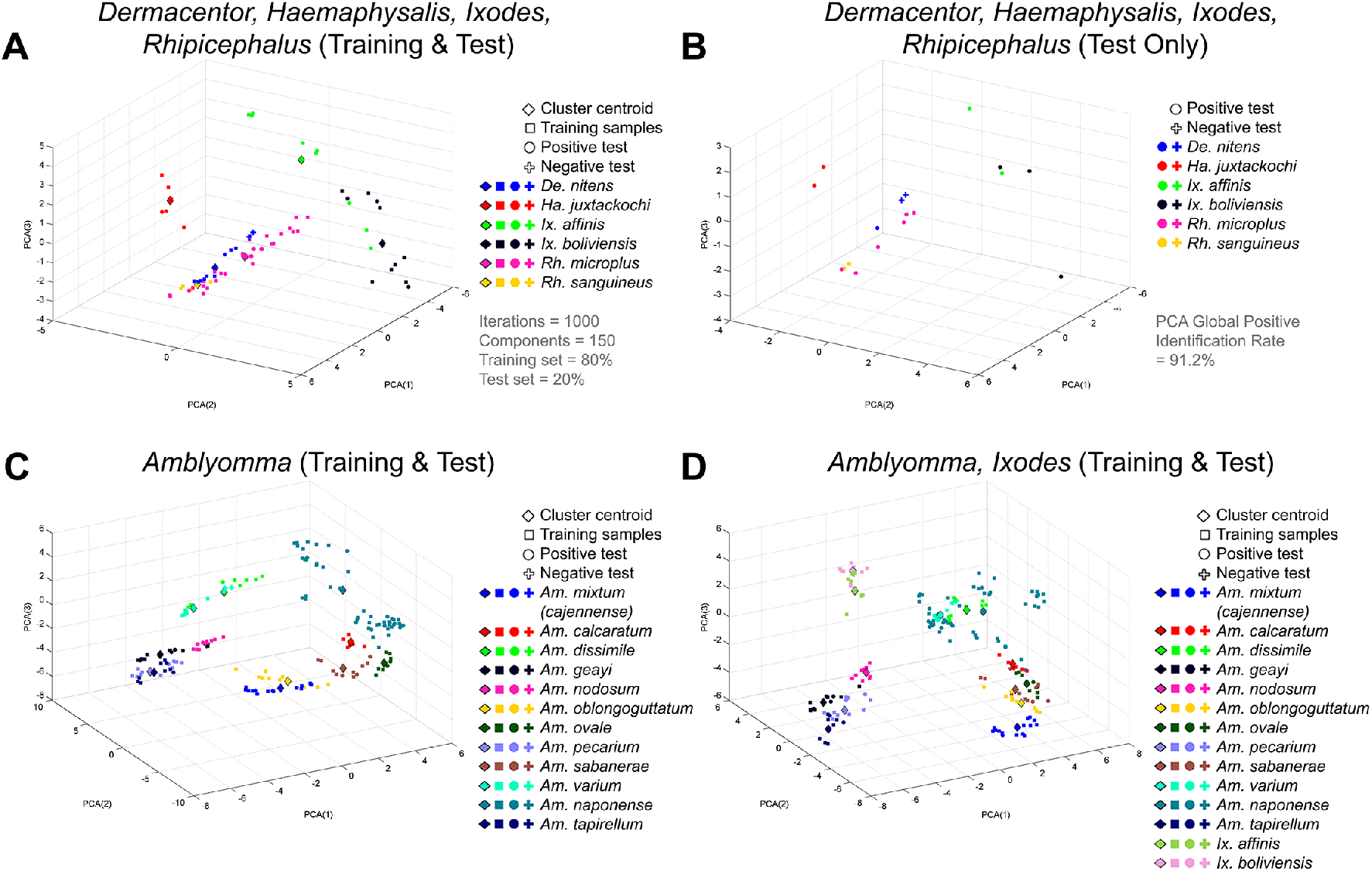
Principal component analysis (PCA) of individual species plotted against first, second and third principal components (PC). All species were classified using a Monte Carlo simulation with 1000 iterations, in which 80% of the samples were used as training set (□) and the remaining 20% as test set (• for positive identifications and + for negative ones). The cluster centroid of each species is also presented in the graph (à). The plots show (A) the training and test sets for the species belonging to the *Dermacentor, Haemaphysalis, Ixodes* and *Rhipicephalus* genera, and (B) only the test sets for better visualization; as well as the training set and test set of (C) *Amblyomma* species alone or (D) *Amblyomma* in combination with *Ixodes* genera. The unsupervised PCA algorithm had a global positive identification rate of 91.2%. These 3D plots represent only one of the 1000 Monte Carlo iterations performed with the algorithm.

In addition, the LDA clustering analysis showed a global positive identification rate of

94.2% (Fig 4; Table 2), with 14 out of 18 species having higher than 97.8 % positive identification rate. The range of positive identification rates went from 100% (best score possible) for *A. mixtum (cajennense), A. nodosum, A. oblongoguttatum, A. ovale, A. varium, A. naponense and R. sanguineus* to 45.6% for *D. nitens*. The 3D representation plots of the LDA clustering displayed that the separation between species was more pronounced than with PCA when comparing species from different genera, confirming the improved quantitative results of the performance of the LDA algorithm (Table 2).

**Figure 4.**
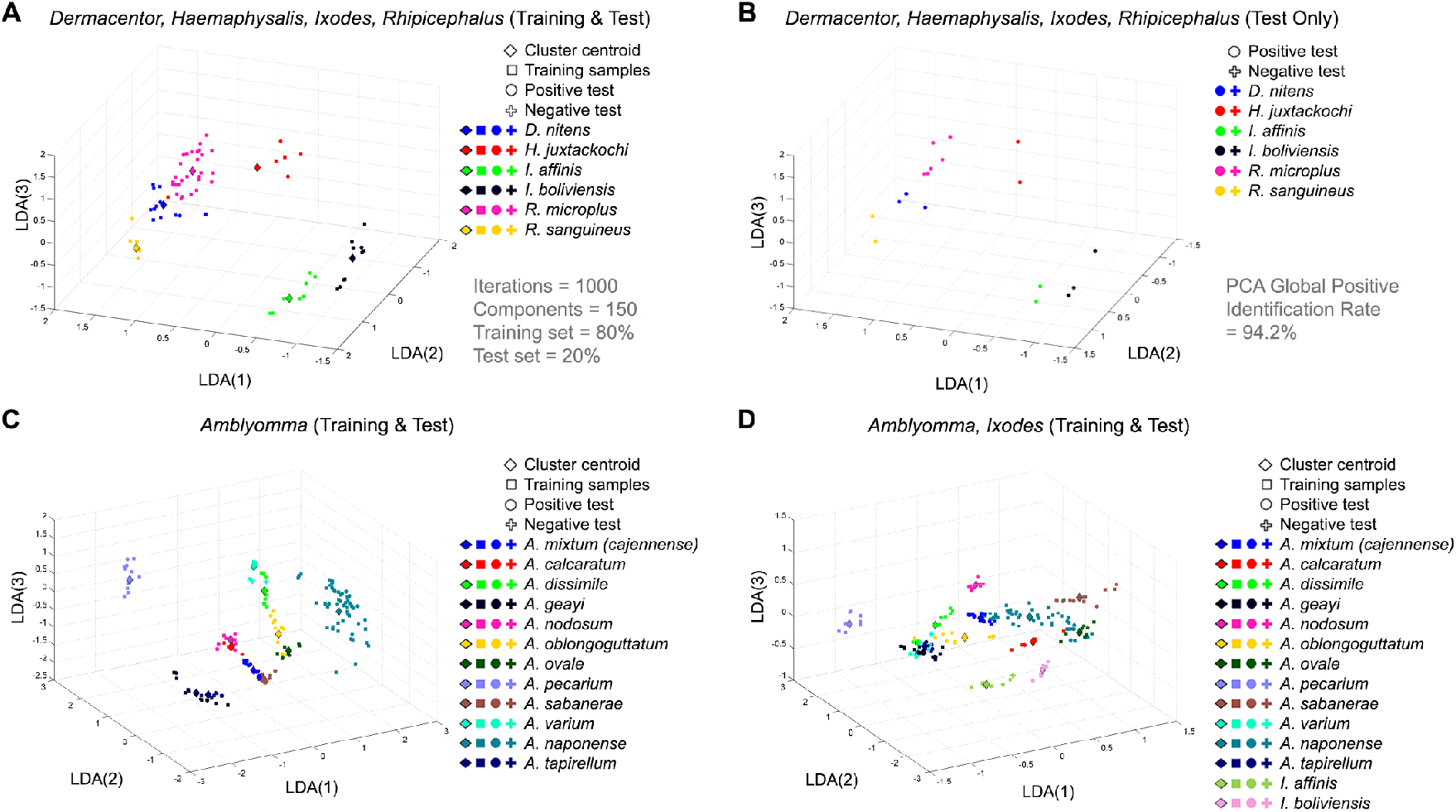
Linear Discriminant Analysis (LDA) applied to spectra from tick species of the genera *Amblyomma, Dermacentor, Haemaphysalis, Ixodes* and *Rhipicephalus*. The plots show (A) the training and test sets for species in the *Dermacentor, Haemaphysalis, Ixodes* and *Rhipicephalus* genera projected over the first three components of the LDA, as well as (B) only the test set for better visualization; and also the training and test sets for (C) the *Amblyomma* genus alone, as well as (D) the *Amblyomma* genus compared to the *Ixodes* genus. These 3D plots represent only one of the 1000 Monte Carlo iterations performed with the algorithm. The supervised LDA algorithm had a 94.2% global positive identification rate.

## Discussion

Our results show that MALDI mass spectra of highly abundant proteins in arthropod legs served as fingerprints to identify samples of 18 species of Neotropical hard ticks using machine learning and pattern recognition algorithms to create a self-curated reference library. We compared smoothed and baseline-corrected spectra generated from unknown field-collected tick samples against the mean spectra from a subset of the same field samples that had already been identified through traditional means. To systematize this process, we used PCA and LDA algorithms to classify mass spectra without prior establishment of a high-quality reference library, which typically requires laboratory-reared specimens that may not be possible to obtain for all species. Global positive identification rates of up to 94.2% were achieved with this methodology, offering a rapid, reliable and objective approach to identify hard tick species, which will likely improve as more specimens are evaluated and included in our database.

These outcomes agree with our previous work [26] in which we used a similar approach to classify field-collected samples of 11 morphologically-identified species of *Anopheles* mosquitoes. In that study, Neotropical *Anopheles* samples were stored dry in silica gel at −20 °C, which seemed to avoid sample degradation and maintain spectral quality. This contrasts with the present study, where most of our specimens were stored in ethanol at −20 °C for several years. Thus, our findings confirm that our novel analytical approach using MALDI and PCA/LDA clustering algorithms is robust for species classification regardless of the arthropod assemblage, sample storing conditions, and the lack of a high-quality reference library. Our results herein also show that both classification algorithms, PCA and LDA, were capable of clustering and recognizing spectra from up to 18 different tick species, including roughly 50 % of Ixodid taxa (e.g., both ecologically dominant and rare taxa) reported for Panama [26, 41].

LDA outcomes were more discriminant and robust than PCA overall, but PCA also classified species from different genera with over 91 % accuracy and consistency. LDA was able to cluster each of the 18 species of ticks with validation and cross-validation scores above 94 %, both between and within genera. As expected, the clustering algorithm was most accurate for distinctly related phylogenetic species (i.e., *Ixodes, Rhipicephalus* and *Haemaphysalis* genera), with higher than 97 % success rate in most of these cases, than for closely related species (i.e., *Amblyomma* genus).

Although the number of samples analyzed for some ixodid species was relatively low, several of these taxa are considered cryptic species complexes [42] and have been implicated as vectors of human pathogens in Panama as well as more broadly, including *A. mixtum* (*cajennense*) and *D. nitens*, the likely vectors of *Rickettsia rickettsii*, known to cause Rocky Mountain spotted fever [43]. We also included samples of *A. tapirellum, A. oblongoguttatum* and *H. juxtakochi*, three species from which human pathogens have been previously isolated [44], such as: *Coxiella*, whose members cause Q fever; *Ehrlichia*, which causes ehrlichiosis infection; and *Rickettsia*, which causes a variety of bacterial infections in humans and other animals. These results are important because our species identification platform can serve as an additional tool for Health Ministries in Panama and other countries, to monitor, predict and manage tick-borne zoonotic pathogens.

Morphological taxonomic identification of ixodid ticks can be enhanced by molecular techniques such as the DNA barcoding [8, 45], but this procedure is laborious, expensive and needs a well-trained lab-technician. Studies show that typical DNA barcoding costs can range from $2 to $5 per sample, with difficult-to-extract samples increasing the cost two-fold or more [46, 47]; while costs associated to MALDI species identification have been calculated to be less than $0.50 per sample [48–50]. Furthermore, a comprehensive repository of DNA sequences (e.g., DNA barcodes) is needed in order to test species limits, yet only a handful of Neotropical tick species are represented in Genbank [51] or BOLD [52] repositories, which could limit identification to the most common taxa only. In addition, DNA barcoding occasionally fails to delimit species boundaries due to ambiguous evolutionary relationships among closely related tick species [45].

The long-term goal of our analytical approach with MALDI is to offer an open-source, web-based platform where users can upload the protein mass spectra of their known and unknown specimens to increase the number of species covered and to improve the power of our clustering algorithms. This crowd-sourced approach could be more cost effective, given that it is not necessary to generate a reference library of well-curated samples. Instead, field samples can be taxonomically assigned as they arrive to the laboratory using a correctly matched protein fingerprint, while unidentified samples can be identified with traditional methods and added as new entries into the growing self-curated reference database.

In conclusion, the present study used MALDI mass spectrometry as a tool to rapidly identify Neotropical specimens of adult hard ticks that had been preserved in ethanol for several years. Our algorithms were capable of identifying specimens from the 18 tick species evaluated, based on their protein spectra “fingerprint” with up to 94% cross-validation capability. This is the first report of the protein mass spectra from the leg for most of these Neotropical tick species. Large arthropod groups such as ticks are difficult to identify with currently available strategies from commercial vendors, forcing the user to lower the “quality” bar of a positive match to enhance the percentage of correct identification. Our MALDI/self-curated library approach, although still serving as an auxiliary technique to traditional identification methods (and not necessarily replacing them), would reduce considerably the number of samples that would require morphological identification or DNA barcoding. This will reduce the time and cost needed to integrate these techniques in routine surveillance programs in Neotropical regions where tick diversity remains relatively uncharacterized.

## Acknowledgements

Special thanks to Philip Davis and Eric Rodríguez from the University of Panama, for assisting with tick collections during the study. We are grateful to Aishwarya Sunderrajan, from the Madras Institute of Technology, India, for her support editing some of the figures for this work. We want to thank the governmental personnel at the Panamanian Ministry of Environment (*MiAmbiente*) for supporting scientific collecting of ticks in Panama.

## Supporting information

### Supporting figures and tables

**S1 Figure.**
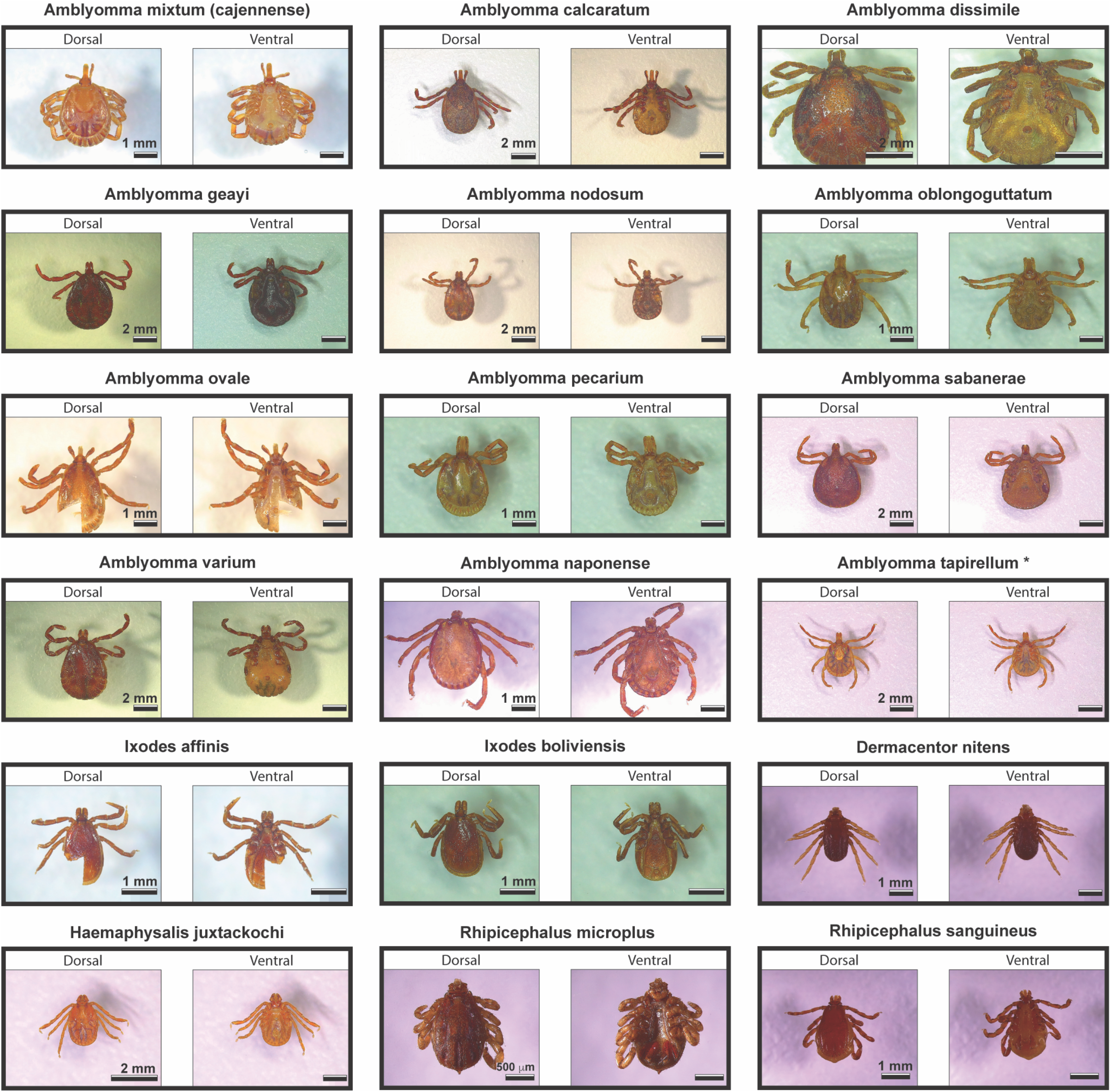
Optical micrographs of Neotropical hard ticks. The image shows the dorsal and ventral sides for all 18 species of hard ticks in the genus *Amblyomma, Dermacentor, Haemaphysalis, Ixodes*, and *Rhipicephalus* used to generate protein spectra with our MALDI mass spectrometry approach.

**S1 Table. Metadata of specimens and species of hard tick (e.g., Ixodidae) collected in Panama. Available at: https://github.com/mjmillerlab/maldi_ticks**

## List of abbreviations

MALDI: matrix-assisted laser desorption/ionization
PCA: principal component analysis
LDA: linear discriminant analysis
DNA: deoxyribonucleic acid
INDICASAT: Institute for Scientific Research and High Technology Services
STRI: Smithsonian Tropical Research Institute
SNI: National System of Investigation
UTP: Technological University of Panama
TOF: time-of-flight
MiAmbiente: Ministry of Environment.

## Ethics approval and consent to participate (Ethics statement)

Not applicable

## Consent for publication

Not applicable

## Availability of data and material

The datasets used and/or analyzed during the current study are available from the corresponding author on reasonable request.

## Authors’ contributions

JRL and RAG designed and developed the experiments. JRL, EA and HJE collected and identified the ticks. AA, RH and MD performed the tests with the MALDI. JRL, JSG, FM, JK and RG analyzed the data and produced the graphs. JRL and RAG wrote the first draft of the paper and EA, LM, KLB, JSG, FM, JK, MJM, HJE, RH, MD, and LFL contributed comments to subsequent versions on it. All authors read and approved the final manuscript.

